# Variability and Impact of Musculoskeletal Modeling Parameters for the Human Elbow

**DOI:** 10.1101/2022.10.29.514351

**Authors:** Russell Hardesty, Byeongchan Jeong, Darren E. Gemoets

**Affiliations:** National Center for Adaptive Neurotechnologies; Department of Veterans Affairs, Stratton Medical Center, Albany, NY; Purdue University

## Abstract

Musculoskeletal modeling has significant potential as a translational and clinical research tool for examining neuromuscular injuries and disorders. However its adoption has been limited due, in part, to the difficulty of measuring the subject-specific physiological measures that define model parameters. These measurements may require substantial time and expensive methods, such as MRI, to determine the parameters of a model and thus ensure its accuracy. We used a Monte Carlo simulation to examine the impact of parameter variability on the ill-defined, inverse approximation of muscle activity. We first amalgamated previously published measurements of the physiological characteristics of the upper/lower arm and the biceps/triceps muscles. We then used the observed distributions of these measurements to set physiologically plausible boundaries on uniform distributions and then generated perturbed parameter sets. We computed the root mean squared error (RMSE) between muscle activity patterns generated by the perturbed model parameters to those generated by the original parameters. Regression models were fit to the RMSE of the approximated muscle activity patterns to determine the sensitivity of the simulation results to variation in each parameter. We found that variation in parameters associated with muscle physiology had the most effect on RMSE, suggesting that these parameters may require subject-specific scaling, whereas parameters associated with skeletal bodies had less effect, and might be safely approximated by their population means.

## INTRODUCTION

Musculoskeletal (MS) modeling is a useful, and increasingly popular, tool for studying the underlying dynamics of movement (Hicks et al., 2015). MS models provide mathematical approximations of the body’s mechanics to relate activity of the nervous system, i.e., the controller, to the behavior of the musculoskeletal system, i.e., the plant. These models have been used to investigate injury prevention/treatment (Marra et al., 2015), design prostheses/orthoses (Sartori et al., 2018), and infer motor control strategies (Al Borno et al., 2020).

The accuracy of these model approximations is dependent, at least in part, on the accuracy and precision of the parameters which describe the model components. For body segments, these parameters include physiological measures such as center-of-mass and inertia; for muscles, they include muscle lengths, pennation angles, etc. The emergence of accessible and standardized platforms, such as OpenSim and MuJoCo, have facilitated the development and dissemination of increasingly complex models, i.e. larger numbers of components and thus parameters. Parameter value selection is commonly addressed by: 1) averaging anthropometric measurements to create a generic model; 2) linear scaling of parameters to subject dimensions; or 3) by creating subject-specific models. Generic models do not account for the inherent inter-subject variability in these parameters, and scaling of these parameters has limited accuracy (Correa et al., 2011; Scheys et al., 2008). Because of these limitations, subject-specific models have been suggested as the preferred approach. However, subject-specific models have their own limitations. For example, some parameters may be difficult, time-consuming, or expensive to approximate and/or measure, such as those requiring MRI imaging. Furthermore, certain applications may benefit from a more generalizable biomechanical description, e.g. biomimetic controllers for prosthetics (Sartori et al., 2018). Therefore, better understanding of both the inherent biological variability of these parameters and their relative contribution in MS simulations is needed to improve modeling assumptions and model design.

Previous studies have attempted to address the impact of parameter uncertainty on simulation outcomes. Many have focused on evaluating the reliability of subject-specific parameters and have generally found them to be robust to measurement errors (Myers et al., 2015; Valente et al., 2014; Hannah et al., 2017). Thus, subject-specific models are generally the preferred approach. Conversely, studies that have evaluated generic or scaled models have shown them to be less reliable (Correa et al., 2011; Nolte et al., 2016; Scheys et al., 2008). However generic models, if generalizable, have several advantages. First, they do not require costly and time-consuming measurements of each subject. Second, they provide a practicable tool for designing human-interface devices that target a diverse population. Third, generic models may capture meaningful relationships that do not greatly vary across individuals (Gritsenko et al., 2016; Hardesty et al., 2020). Furthermore, parameter variability is not entirely unconstrained. The physiological measures that these parameters represent may be co-dependent, e.g., segment mass and inertia. Or they may be irrelevant, i.e., they do not significantly influence simulation results. These factors can decrease the number of subject-specific parameters that must be measured. In fact, in at least one study, anatomical variability was found to follow multimodal distributions, suggesting that inter-subject variability could be taken into account by using a finite set of models (Santos and Valero-Cuevas, 2006). Constraining the biologically plausible parameter space could enable robust model development while limiting the time and cost associated with creating subject-specific models.

Here, we investigate the impact of variability in musculoskeletal parameters relevant to modeling motion about the elbow joint. We ask three questions: First, what is the inherent variability of each of these parameters? Second, do any of these parameters correlated with one another such that a subset of parameters could accurately predict all of them? Third, for which parameters does their variability have the greatest effect on the outcome of musculoskeletal simulations? To answer these questions, we amalgamated anthropometric measurements from previously published studies to create a database of parameters for the upper and lower arms and the biceps and triceps muscle. We then determined the variability of these parameters and their interdependence. Finally, beginning with the observed parameter distributions, we use a Monte Carlo simulation and multiple regression models to determine the sensitivity of inverse simulations to the variability in each of these parameters.

## METHODS

### Musculoskeletal Model

We created a 1 degree-of-freedom (DOF) model of the elbow joint using the OpenSim v4.3 musculoskeletal modeling software (https://simtk.org/projects/opensim). The model comprises two rigid bodies representing the upper and lower arm joined by a single revolute joint representing the elbow. The initial parameter quantities, kinematic descriptions of the elbow joint, and muscle geometry were adapted from the MOBL-ARMS Dynamic Upper Limb (Saul et al., 2015a; McFarland et al., 2019). Muscle geometry definitions were updated to reference this two-body system such that the muscle lengths, tendon lengths, and muscle moment arms were consistent with the MOBL arm across the full range of motion of the simplified model (see Figure 1). The model was created using custom Python code (v3.8) (Van Rossum and Drake Jr, 1995) and the OpenSim API (Delp et al., 2007; Seth et al., 2018).

**Figure 1.**
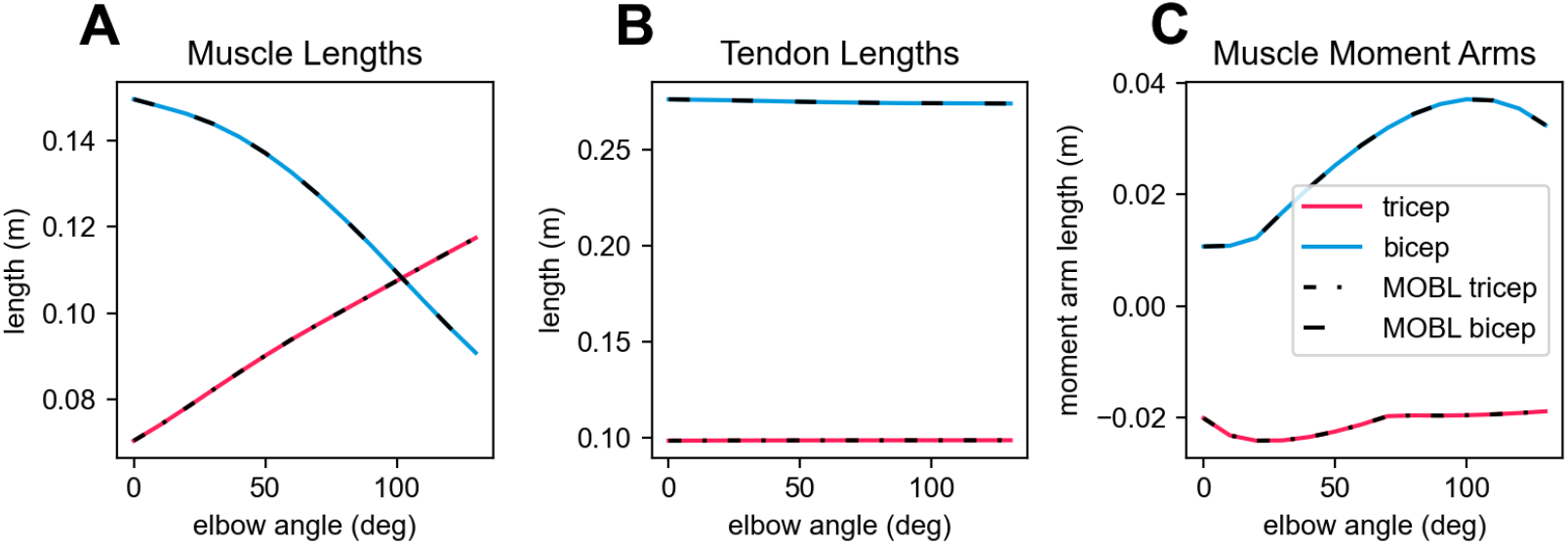
Musculoskeletal dynamics of simplified elbow model are consistent with the MOBL Dynamics Arm model. Muscle lengths (A), tendon lengths (B), and muscle moment arms (C) are shown across the full range of the elbow joint.

### Data Collection

To approximate the inherent biological variability of model parameters, we reviewed anthropometric publications on the upper arm and forearm to extract measurements of physiological characteristics used in musculoskeletal model parameters. We defined a global Cartesian coordinate system where the x-direction was along the posterior-anterior axis, the y-direction along the superior-inferior axis, and the z-direction along the lateral-distal axis, as shown in Figure 2. Local coordinates for the upper arm and forearm where defined with origins at the center of the shoulder and elbow joints respectively. The coordinates were defined such that during neutral posture, the arm was orthogonal to the ground and all local coordinates were equal to 0. Published parameters that were defined relative to differing coordinate systems were transformed so that all values were relative to this predefined coordinate system. Reported measurements were coalesced into an SQL database to perform statistical analysis and subsequent computational simulations. This database and the relevant Python code are available at https://github.com/NeuroEng/ms_db-git.

**Figure 2.**
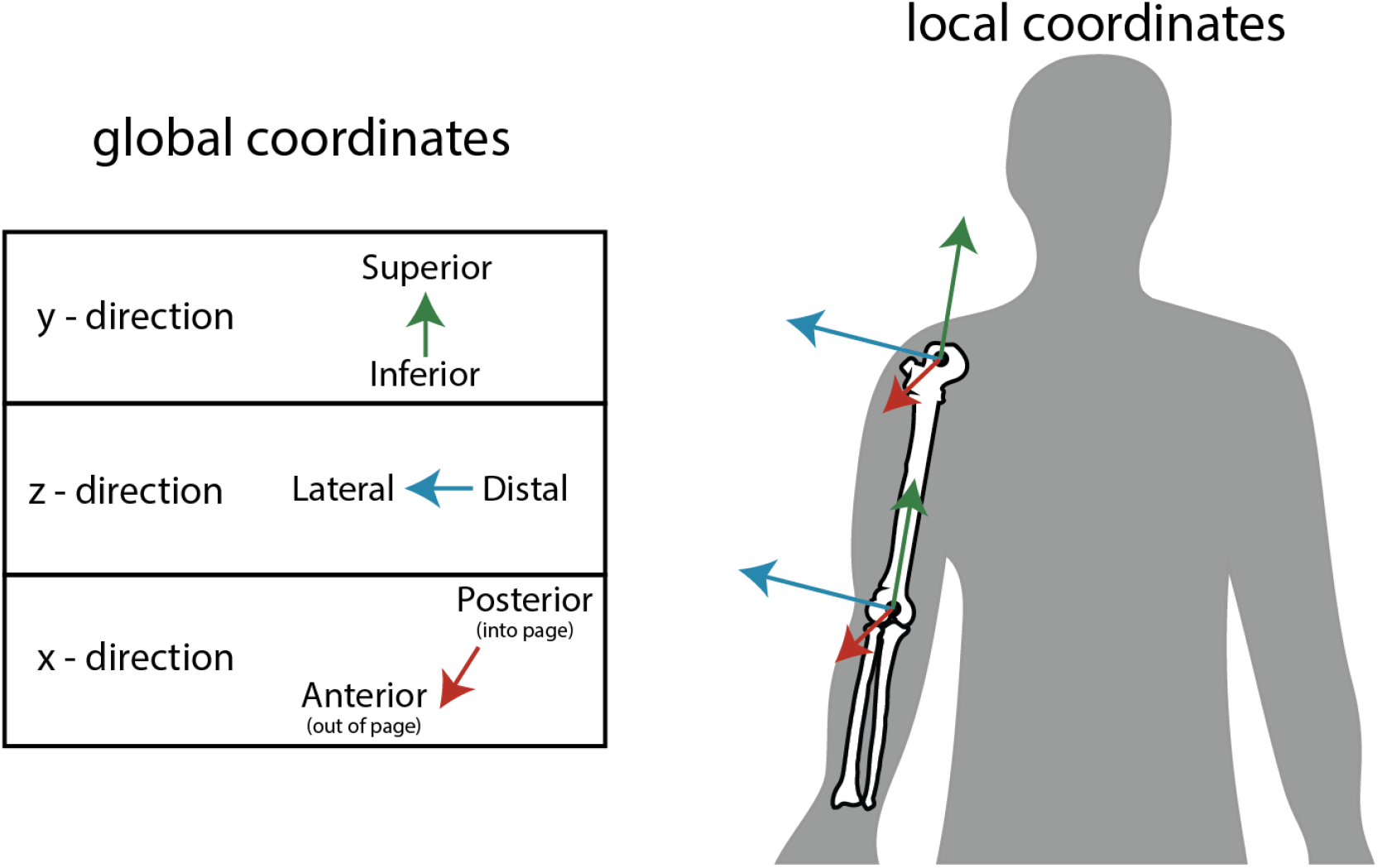
The global and local Euler coordinate definitions for all parameter values are shown. The global coordinate was defined such that at neutral posture (all joint angles equal to 0) the arm is parallel to gravity. Local coordinate origins were located in the center of the shoulder and elbow joint.

### Biological variability

To determine the inherent parameter variability, we calculated summary statistics including the mean, standard deviation, median, interquartile range, coefficient of variations, and confidence intervals for each parameter in the amalgamated dataset. These summary statistics were then used to constrain our coefficient exploration (see Computational Simulations). We then performed a linear regression between parameters that are likely to be interdependent, e.g. mass vs. inertia for bodies and muscles and calculated the correlation coefficient. This determined whether all parameters could be accurately predicted from a subset of parameters. R (R Core Team, 2020) or Python (Van Rossum and Drake Jr, 1995) were used for the sensitivity analyses and all statistical analyses.

### Computational Simulations

We chose to evaluate the sensitivity of musculoskeletal parameters in inverse simulations of the muscle activity that produces a predefined desired movement. Inverse musculoskeletal simulations are ill-defined, i.e., they may be satisfied with multiple solutions, a feature of musculoskeletal physiology termed “motor redundancy.” When there is no explicit solution, this problem can be addressed by various optimization methods. Because this problem is ill-defined, we found that these optimization procedures would be particularly susceptible to variability in model parameters. We used OpenSim’s computed muscle control (CMC) optimization procedure to calculate muscle activity patterns capable of generating a sigmoidal flexion of the elbow joint. The CMC procedure combines proportional-derivative control with a static optimization to optimize muscle activity over a designated time window (10ms) while minimizing total activation (*a*)

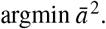

The feedback gain was 100 for position error (*K*_*p*_) and 20 for velocity error (*K*_*v*_). A more thorough description of the CMC procedure can be found in (Thelen and Anderson, 2006). The desired movement was a physiologically-realistic sigmoidal flexion of the elbow joint from 0 to 120 degrees over 1 sec with a 0.5 sec hold both before and after the movement to ensure that the simulations achieved equilibrium.

To evaluate parameter sensitivity, we performed a Monte Carlo procedure where each parameter was randomly assigned from a uniform distribution bounded by the 95% confidence interval for the mean calculated from the amalgamated literature values. The CMC optimization was repeated for each set of parameter values, as shown in Figure 3, and the resultant muscle activation profiles were compared with results obtained the unperturbed parameter values used in the MOBL model. The root-mean-square error (RMSE) between the original and perturbed muscle activities was calculated.

**Figure 3.**
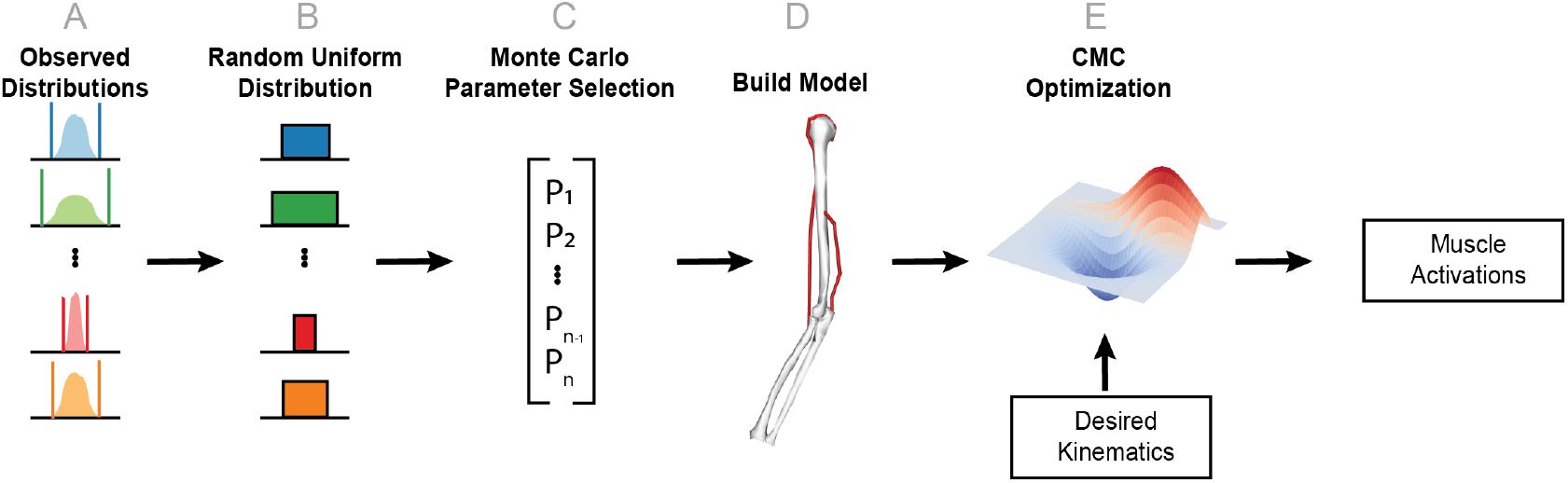
A schematic representation of the simulation procedure is shown. This procedure is repeated for each simulation (N=1000). Parameter distributions were approximated from an amalgamated data set of previously reported measurements (A). These distributions were then used to create a random uniform distribution constrained by the 95-percentile confidence interval (B). A particular parameter set is generated using a Monte Carlo re-sampling of the uniform distributions (C). An OpenSim model of the elbow is then generated using this parameter set (D). Finally, the CMC optimization is performed using predefined desired kinematics and the newly generated model (E).

### Sensitivity Analysis

We used multiple linear regression models to assess the impact of each parameter on the simulation results. The data were standardized (i.e., each variable was centered on its mean and scaled by its standard deviation). Next, a least squares multiple regression model was fit to the standardized data with the RMSE values as the response variable and the sampled MS parameter values as the predictor variables (see, for example, Saltelli et al. (2008) for more details). This standardization allowed for across-parameter comparisons, and each regression parameter estimate can be interpreted as standardized change in RMSE per standardized change in MS parameter. Main effects models are reported in the Results section. Models with two-way interactions – which explore the joint effect of parameters on RMSE – are reported in the Supplemental materials.

## RESULTS

### Parameter Variability

We first assessed the distribution of musculoskeletal modeling parameters as reported in the literature (see Figures 4 and 5). We calculated the mean, standard deviation, median, interquartile range (IQR), the coefficient of variation, and 95% confidence intervals for each parameter. These summary statistics for the upper arm, forearm, biceps, and triceps are shown in tables 5, 6, 7, and 8, respectively. Amongst body parameters, the center-of-mass, in the x (anterior-posterior) and in the z (medial-lateral) direction were the most variable, as shown by their coefficients of variation (center-of-mass (x): 3.1 and -1.7; center-of-mass (z): -2.2 and 10.7 for the upper and lower arm, respectively). In contrast, muscle parameters all had coefficients of variation less than 1 (range: 0.12 - 0.49) demonstrating that muscle parameter magnitudes are less variable relative to their mean value.

**Table 1.** Number of individual measurements obtained from each reference for the humerus.

| reference | mass | com (x) | com (y) | com (z) | Ixx | Iyy | Izz |
| --- | --- | --- | --- | --- | --- | --- | --- |
| Chandler et al. (1975) | 12 | 12 | 12 | 12 | 12 | 12 | 12 |
| Ho et al. (2013) | - | - | - | - | 1 | 1 | 1 |
| Jensen (1978) | 3 | - | - | - | 3 | 3 | 3 |
| McConville et al. (1980) | - | 2 | 2 | 2 | 2 | 2 | 2 |
| Nikolova (2010) | - | - | - | - | 2 | 2 | 2 |
| Nikolova and Toshev (2007) | 2 | - | - | - | 2 | 2 | 2 |
| Saul et al. (2015b) | 1 | 1 | 1 | 1 | 1 | 1 | 1 |
| Shan and Bohn (2003) | - | - | - | - | 2 | 2 | 2 |
| Veeger et al. (1991) | 7 | - | 7 | - | 7 | - | 7 |
| Veeger et al. (1997) | 4 | - | 4 | - | 4 | - | 4 |
| Young et al. (1983) | - | 2 | 2 | 2 | 2 | 2 | 2 |

**Table 2.** Number of individual measurements obtained from each reference for the forearm.

| reference | mass | com (x) | com (y) | com (z) | Ixx | Iyy | Izz |
| --- | --- | --- | --- | --- | --- | --- | --- |
| Chandler et al. (1975) | 12 | 12 | 12 | 12 | 12 | 12 | 12 |
| Ho et al. (2013) | - | - | - | - | 1 | 1 | 1 |
| Jensen (1978) | 3 | - | - | - | 3 | 3 | 3 |
| Jensen and Fletcher (1993) | - | - | - | - | 2 | 2 | 2 |
| McConville et al. (1980) | - | 2 | 2 | 2 | 2 | 2 | 2 |
| Nikolova (2010) | - | - | - | - | 2 | 2 | 2 |
| Nikolova and Toshev (2007) | 2 | - | - | - | 2 | 2 | 2 |
| Shan and Bohn (2003) | - | - | - | - | 2 | 2 | 2 |
| Veeger et al. (1991) | 7 | - | 7 | - | 7 | - | 7 |
| Veeger et al. (1997) | 4 | - | 4 | - | 4 | - | 4 |
| Young et al. (1983) | - | 2 | 2 | 2 | 2 | 2 | 2 |

**Table 3.** Number of individual measurements obtained from each reference for the biceps muscle.

| reference | max iso. force | opt. fiber length | tendon slack length | pen. angle |
| --- | --- | --- | --- | --- |
| Amis et al. (1979) | - | - | - | - |
| An et al. (1981) | - | - | - | - |
| Garner and Pandy (2001) | 1 | 1 | 1 | 1 |
| Holzbaaur et al. (2005)5 | 1 | 1 | 1 | - |
| Koo (2001) | - | 5 | 5 | - |
| Langenderfer et al. (2004a) | - | 1 | 1 | - |
| Murray et al. (2000) | - | 1 | 1 | 1 |
| Peterson and Rayan (2011) | - | 1 | - | 1 |
| Saul et al. (2015b) | 2 | 2 | 2 | - |
| Veeger et al. (1997) | - | - | - | 1 |
| Winters and Stark (1988) | - | - | 1 | 1 |

**Table 4.** Number of individual measurements obtained from each reference for the triceps muscle.

| reference | max iso. force | opt. fiber length | tendon slack length | pen. angle |
| --- | --- | --- | --- | --- |
| Amis et al. (1979) | - | - | - | - |
| Amis et al. (1979) | - | - | - | - |
| Garner and Pandy (2001) | 1 | 1 | 1 | 1 |
| Holzbaaur et al. (2005) | 1 | 1 | 1 | 1 |
| Koo (2001) | - | 5 | 5 | - |
| Langenderfer et al. (2004b) | - | 1 | 1 | 1 |
| Murray et al. (2000) | - | 1 | 1 | 1 |
| Peterson and Rayan (2011) | - | 1 | - | 1 |
| Saul et al. (2015b) | 2 | 2 | 2 | 2 |
| Veeger et al. (1997) | - | - | - | 1 |
| Winters and Stark (1988) | - | - | 1 | 1 |

**Table 5.** Summary statistics for body parameters of the upper arm.

| Parameter | Mean | STD | Median | IQR | Coef Variance | 95% CI for Mean |
| --- | --- | --- | --- | --- | --- | --- |
| mass | 1.78867 | 0.407918 | 1.815 | 0.43757 | 0.228056 | 1.632477 - 1.944873 |
| center-of-mass (x) | 0.00336235 | 0.0104634 | 0.005 | 0.0109 | 3.11192 | -0.001938 - 0.008663 |
| center-of-mass (y) | -0.162539 | 0.039635 | -0.15935 | 0.03575 | -0.24385 | -0.177994 - -0.147083 |
| center-of-mass (z) | -0.0105088 | 0.0227707 | -0.008 | 0.0281 | -2.16682 | -0.022043 - 0.001026 |
| inertia (xx) | 0.0110324 | 0.00530084 | 0.01224 | 0.0069325 | 0.48048 | 0.009267 - 0.012798 |
| inertia (yy) | 0.00551769 | 0.00599471 | 0.0024785 | 0.00659845 | 1.08645 | 0.003136 - 0.0079 |
| inertia (zz) | 0.00894227 | 0.00571488 | 0.0095562 | 0.0105385 | 0.639086 | 0.007039 - 0.010846 |

**Table 6.**
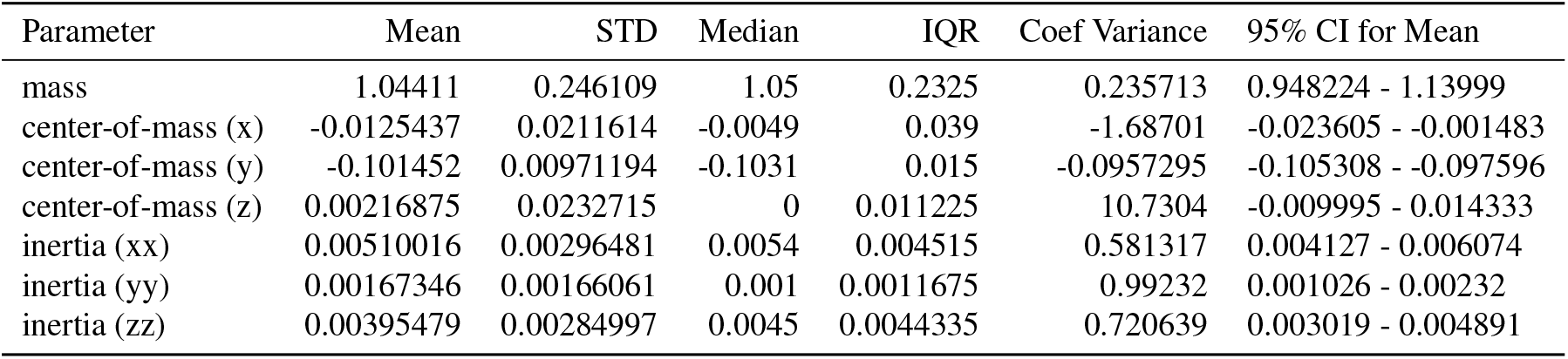
Summary statistics for body parameters of the forearm.

| Parameter | Mean | STD | Median | IQR | Coef Variance | 95% CI for Mean |
| --- | --- | --- | --- | --- | --- | --- |
| mass | 1.04411 | 0.246109 | 1.05 | 0.2325 | 0.235713 | 0.948224 - 1.13999 |
| center-of-mass (x) | -0.0125437 | 0.0211614 | -0.0049 | 0.039 | -1.68701 | -0.023605 - -0.001483 |
| center-of-mass (y) | -0.101452 | 0.00971194 | -0.1031 | 0.015 | -0.0957295 | -0.105308 - -0.097596 |
| center-of-mass (z) | 0.00216875 | 0.0232715 | 0 | 0.011225 | 10.7304 | -0.009995 - 0.014333 |
| inertia (xx) | 0.00510016 | 0.00296481 | 0.0054 | 0.004515 | 0.581317 | 0.004127 - 0.006074 |
| inertia (yy) | 0.00167346 | 0.00166061 | 0.001 | 0.0011675 | 0.99232 | 0.001026 - 0.00232 |
| inertia (zz) | 0.00395479 | 0.00284997 | 0.0045 | 0.0044335 | 0.720639 | 0.003019 - 0.004891 |

**Table 7.** Summary statistics for muscle parameters of the biceps (long head).

| Parameter | Mean | STD | Median | IQR | Coef Variance | 95% CI for Mean |
| --- | --- | --- | --- | --- | --- | --- |
| max iso. force | 516.852 | 82.2234 | 525.1 | 57.8475 | 0.159085 | 414.296121 - 619.408879 |
| opt. length | 0.12411 | 0.0244155 | 0.116 | 0.027575 | 0.196724 | 0.106528 - 0.141692 |
| tendon slack length | 0.249955 | 0.0308595 | 0.261 | 0.0464 | 0.123461 | 0.228872 - 0.271037 |
| pen. angle | 11.6667 | 2.35702 | 10 | 2.5 | 0.202031 | 8.066052 - 15.267281 |

**Table 8.** Summary statistics for muscle parameters of the triceps (lateral head).

| Parameter | Mean | STD | Median | IQR | Coef Variance | 95% CI for Mean |
| --- | --- | --- | --- | --- | --- | --- |
| max iso. force | 832.043 | 255.056 | 717.5 | 161.142 | 0.306543 | 513.913282 - 1150.171718 |
| opt. length | 0.10713 | 0.0322784 | 0.1054 | 0.024 | 0.301301 | 0.083886 - 0.130374 |
| tendon slack length | 0.145718 | 0.037968 | 0.167 | 0.0733 | 0.260558 | 0.11978 - 0.171657 |
| pen. angle | 16.1429 | 7.97189 | 15 | 11.5 | 0.493834 | 9.111915 - 23.173799 |

**Figure 4.**
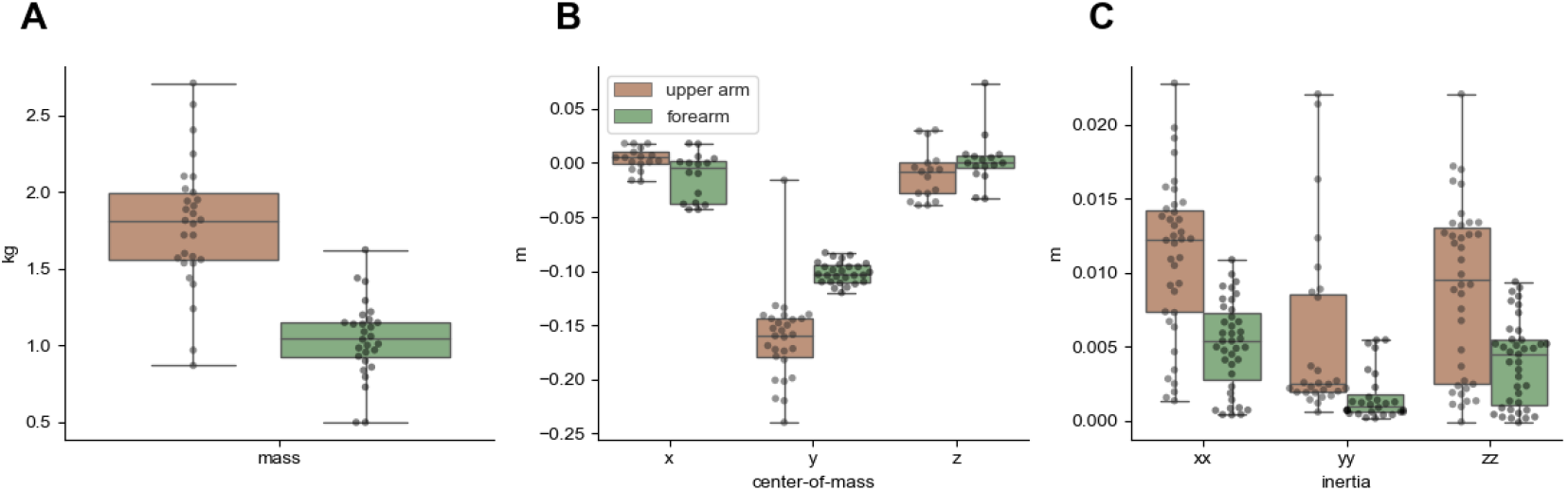
The distributions of body parameters are shown. Dots correspond to individual values obtained from previously published studies.

**Figure 5.**
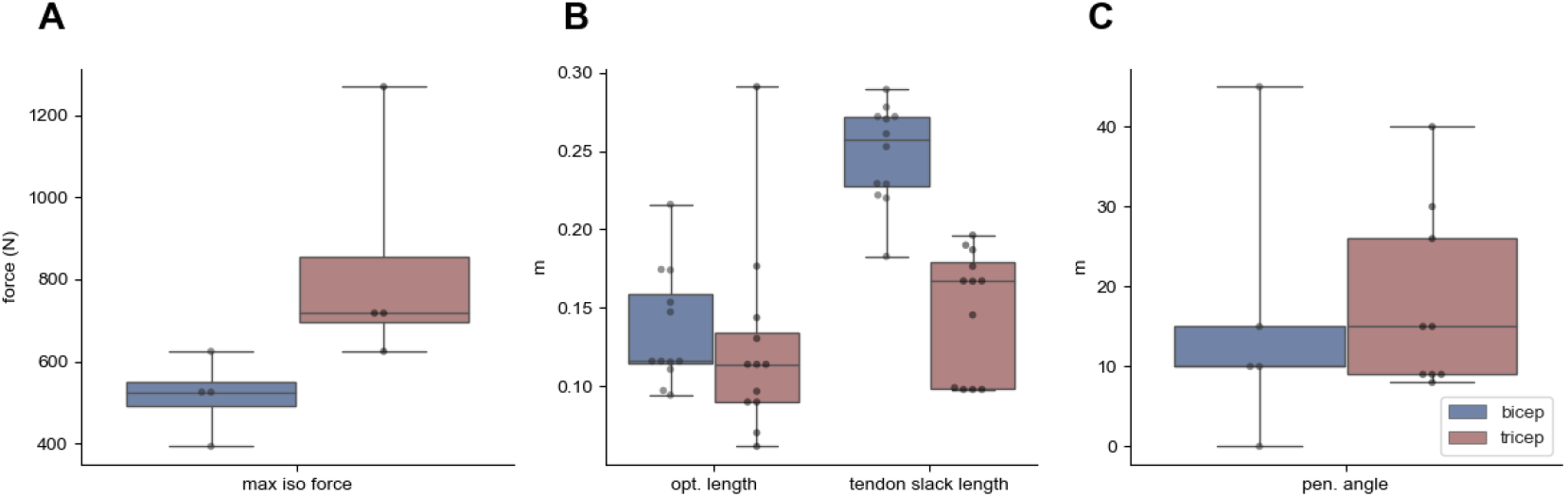
The distributions of muscle parameters are shown. Dots correspond to individual values obtained from previously published studies.

### Parameter Correlation

Next, we calculated the Pearson’s correlation coefficient (*r*) and the variance explained (*r*^2^) to determine whether parameter values may be approximated from a subset of measured values (see Figure 6). Body parameters showed weak to moderate correlations with one another (*r*^2^ range: 0.000317 - 0.521486). The strongest correlation was between inertia (Ixx) and mass and center-of-mass in the y-direction. The center-of-mass in the x- and z-directions, which were the most variable parameters in our data set, were only weakly correlated with other body parameters. The largest *r*^2^ for center-of-mass in the x-direction was 0.04 and the largest in the z-direction was 0.25. Muscle parameters also showed weak correlations, with two exceptions: (1) maximum isometric force and pennation angle (*r*^2^ : 0.69) and (2) maximum isometric force and optimal fiber length (*r*^2^ : 0.88). Overall, these results suggest that body parameters, particularly center-of-mass in the x- and z-directions are not only more variable, but cannot be accurately approximated from linear regression with other body parameter values.

**Figure 6.**
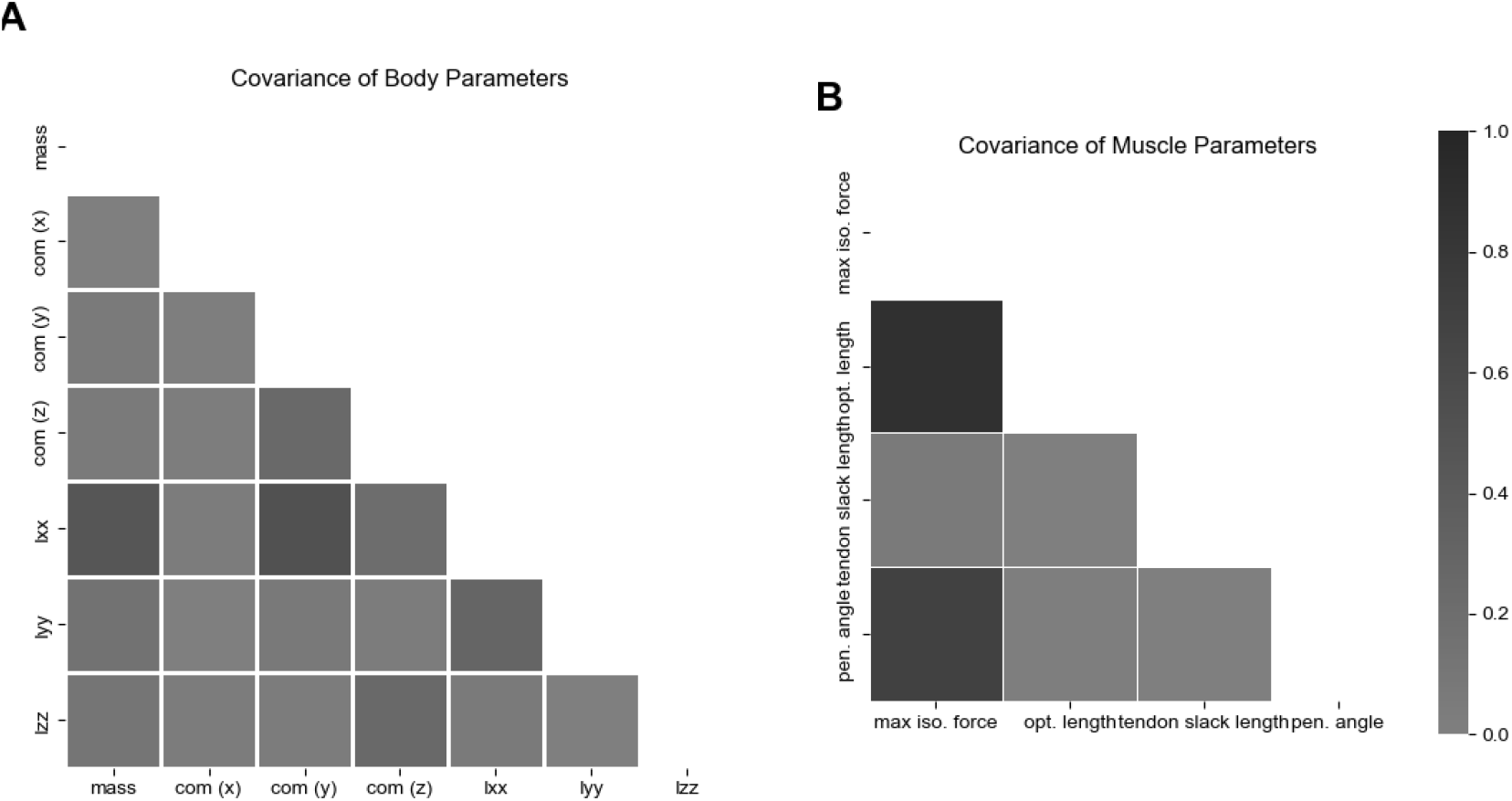
Correlation matrices for body (A) and muscle (B) parameters are shown. The color denotes the variance explained, *r*^2^, for each correlation. The upper diagonal has been removed to avoid redundancy and provide clarity.

### Simulation Results

The CMC optimization successfully computed muscle activations for 990 of the 1000 simulations. For 10 simulations, the optimization was unable to find an acceptable muscle activity profile within the integrator tolerance (0.00001). The desired elbow angle is show in Figure 7A; the movement began with the elbow extended at an angle of 0 radians. The muscle activity profiles for individual simulations are shown as black lines in Figure 7B and C. The mean activity and standard deviation across simulations are shown as solid lines and shaded regions, respectively. As expected, the triceps is activated at the beginning of the simulation; it maintains the elbow in an extended posture prior to movement onset. Once movement begins, triceps activity generally decreases and the biceps activity begins to increase to flex the elbow joint. While this behavior was relatively consistent across the majority of simulations, the timing and extent of triceps deactivation and biceps activation differed depending upon the perturbed parameters. Interestingly, the triceps error varied to a larger extent than that of the biceps (see Figure 8). Because the triceps activity was generally larger than that of the biceps, we computed the coefficients of variation for triceps and biceps error to normalize this error to the magnitude. Even after this normalization triceps profiles varied more than biceps profiles (triceps CV: 0.86; biceps CV: 0.35).

**Figure 7.**
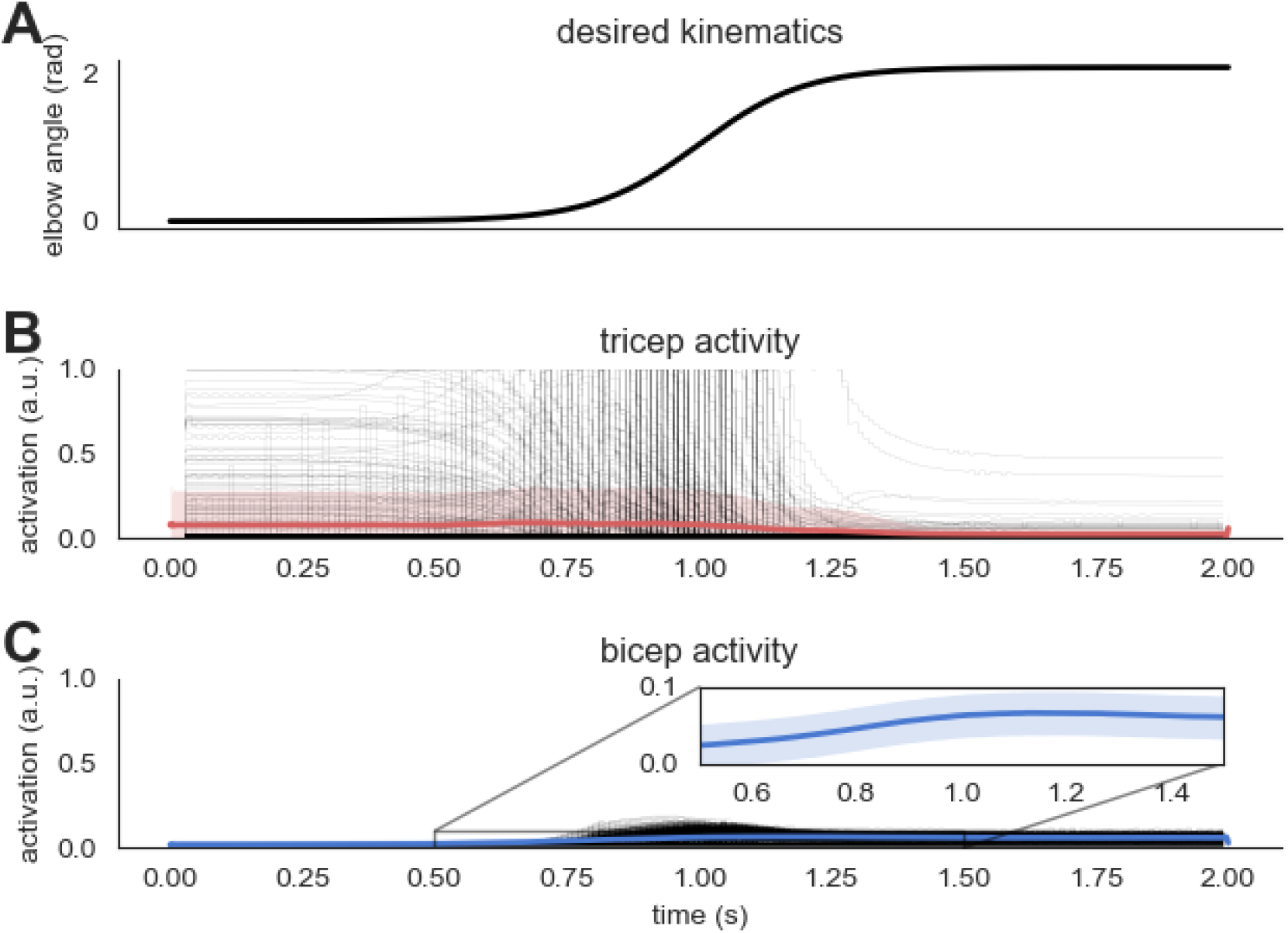
Panel A shows the desired kinematics used for the CMC optimization. Panels B and C show the simulated activity of the triceps and biceps muscles, respectively. Black lines denote individual simulation results. The solid red and blue lines show the average activity across all simulations, while the shaded region shows the standard deviation across simulations.

**Figure 8.**
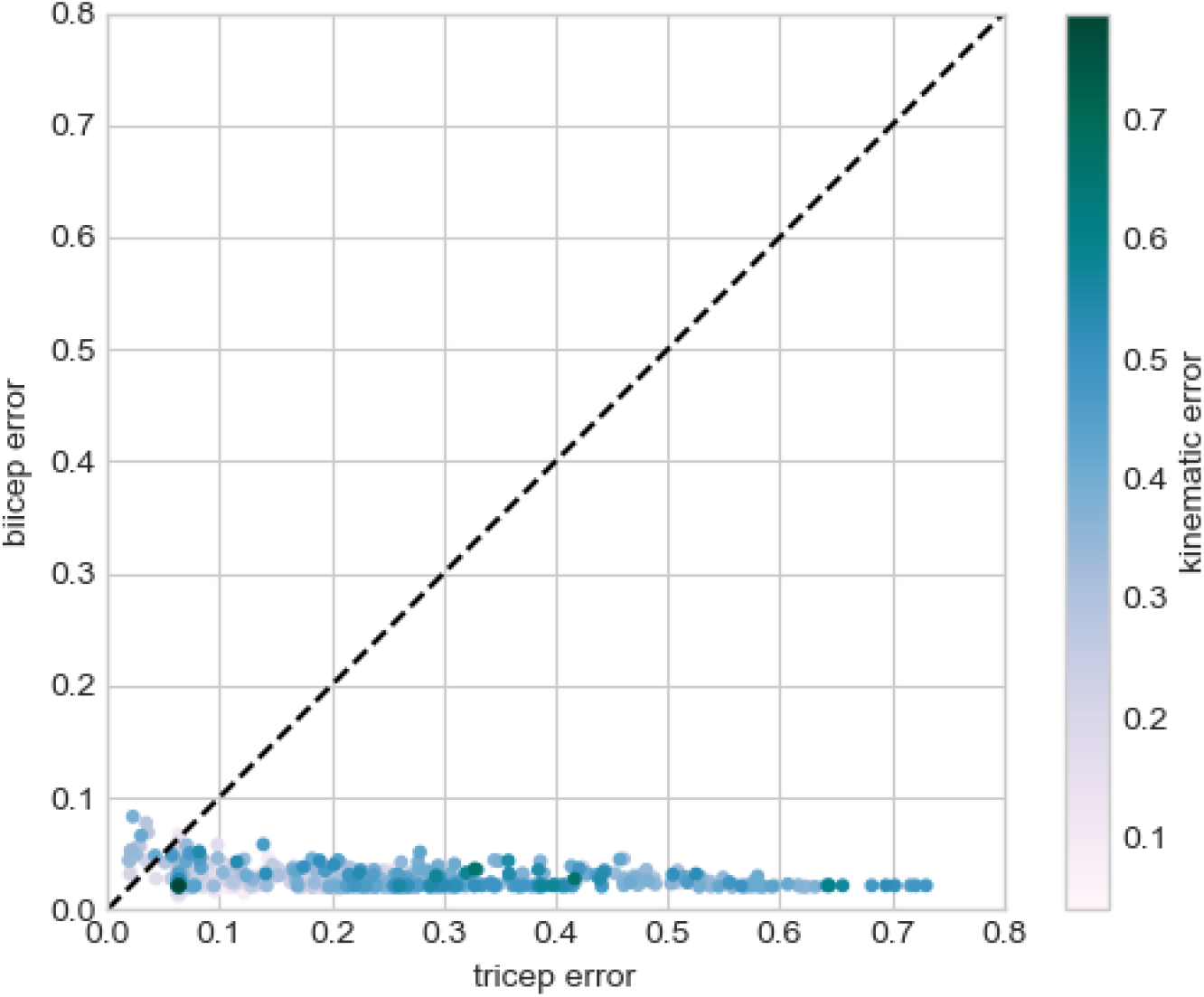
Error was calculated for both the triceps and biceps muscle activity. The error was calculated as the RMSE between the original model parameters and the perturbed parameters for each simulation. Each dot denotes an individual simulation and the color corresponds to the kinematic error. The dashed line is the one-to-one line.

### Sensitivity Analysis

We performed a multiple linear regression of parameter magnitudes to RMSE of both triceps and biceps. RMSE is calculated between the muscle activity patterns generated from our default parameter values and the muscle activity patterns generated from the perturbed parameters in our Monte Carlo simulation.

Table 9 shows the largest regression parameter estimates (in magnitude) with the RMSE for the biceps as the response, with *r*^2^ = 0.70 for this model. Note that positive regression parameters increase RMSE, and because the data are standardized the estimates are comparable. Tendon length and optimal fiber length for both the biceps and triceps have the largest impact in RMSE, along with the biceps maximum force. We expect some false-positive results may occur and suspect that this may be the case for the triceps pennation angle and lower-arm COM (y-direction), as their estimates are the lowest amongst significant parameters. It should be noted that our simulations had 66 duplicate biceps error values. Because this was a small fraction of the total number of simulations (*n* = 1000), we believe that these duplicates may be the result of: 1) sensitive parameters having similar values in this small subset of simulations; 2) a floor effect due to the fact that movement was being generated entirely by passive forces; and/or 3) limited numerical precision. Table 10 shows the largest regression parameter estimates with the RMSE for the triceps as the response (with *r*^2^ = 0.65). Similar to our regression results using the biceps RMSE as the response, the largest regression parameter estimates were for muscle parameters. The largest estimates being the optimal length of the biceps and the tendon slack length of both muscles. There were more repeated error values for the triceps (*n* = 170). Because these were present in a minority of simulations, we included all values in the regression model.

**Table 9.** Regression parameter estimates for biceps RMSE, in order of decreasing value (in magnitude).

|  | Estimate | Std. Error | t value | P-value |
| --- | --- | --- | --- | --- |
| max. force (biceps) | -0.45 | 0.02 | -25.55 | 0.00 |
| tendon slack length (biceps) | 0.42 | 0.02 | 23.96 | 0.00 |
| tendon slack length (triceps) | -0.36 | 0.02 | -20.19 | 0.00 |
| opt. length (biceps) | 0.32 | 0.02 | 17.78 | 0.00 |
| opt. length (triceps) | -0.21 | 0.02 | -11.55 | 0.00 |
| lower arm mass | 0.19 | 0.02 | 10.41 | 0.00 |
| max. force (triceps) | 0.17 | 0.02 | 9.70 | 0.00 |
| lower arm COM (y) | -0.09 | 0.02 | -5.01 | 0.00 |
| pen. angle (triceps) | 0.05 | 0.02 | 2.98 | 0.00 |
| lower arm COM (x) | -0.04 | 0.02 | -2.21 | 0.03 |
| upper arm Iyz | 0.04 | 0.02 | 2.10 | 0.04 |
| upper arm COM (z) | 0.03 | 0.02 | 1.52 | 0.13 |
| upper arm Ixx | 0.02 | 0.02 | 1.36 | 0.17 |
| lower arm Izz | -0.02 | 0.02 | -1.23 | 0.22 |
| lower arm Iyy | 0.02 | 0.02 | 1.13 | 0.26 |
| upper arm Ixy | 0.02 | 0.02 | 1.12 | 0.26 |
| lower arm Ixy | 0.01 | 0.02 | 0.83 | 0.41 |
| lower arm Ixx | 0.01 | 0.02 | 0.66 | 0.51 |
| upper arm COM (x) | -0.01 | 0.02 | -0.63 | 0.53 |
| lower arm COM (z) | -0.01 | 0.02 | -0.51 | 0.61 |

**Table 10.** Regression parameter estimate for triceps RMSE, in order of decreasing value (in magnitude).

|  | Estimate | Std. Error | t value | P-value |
| --- | --- | --- | --- | --- |
| tendon slack length (triceps) | -0.45 | 0.02 | -23.63 | 0.00 |
| opt. length (biceps) | -0.44 | 0.02 | -22.93 | 0.00 |
| tendon slack length (biceps) | -0.39 | 0.02 | -20.30 | 0.00 |
| pen. angle (triceps) | 0.23 | 0.02 | 11.91 | 0.00 |
| opt. length (triceps) | -0.09 | 0.02 | -4.52 | 0.00 |
| max. force (biceps) | 0.09 | 0.02 | 4.51 | 0.00 |
| max. force (triceps) | -0.05 | 0.02 | -2.44 | 0.01 |
| lower arm COM (z) | 0.04 | 0.02 | 2.13 | 0.03 |
| upper arm mass | 0.04 | 0.02 | 1.99 | 0.05 |
| upper arm COM (x) | 0.04 | 0.02 | 1.90 | 0.06 |
| lower arm COM (x) | -0.04 | 0.02 | -1.85 | 0.07 |
| lower arm Iyy | -0.03 | 0.02 | -1.82 | 0.07 |
| upper arm Iyz | -0.03 | 0.02 | -1.74 | 0.08 |
| lower arm Ixz | -0.03 | 0.02 | -1.72 | 0.09 |
| lower arm Ixx | -0.03 | 0.02 | -1.58 | 0.11 |
| lower arm Ixy | -0.03 | 0.02 | -1.51 | 0.13 |
| pen. angle (biceps) | 0.03 | 0.02 | 1.49 | 0.14 |
| upper arm Ixz | -0.03 | 0.02 | -1.45 | 0.15 |
| lower arm Izz | 0.02 | 0.02 | 0.87 | 0.39 |
| upper arm Iyy | 0.02 | 0.02 | 0.84 | 0.40 |

## DISCUSSION

The goal of this study was to evaluate the inherent variability in physiological measures used in musculoskeletal models of the human elbow and to determine the sensitivity of inverse dynamics optimizations to these parameters’ variability. Several other studies have evaluated parameter sensitivity in musculoskeletal models, but these have focused primarily on the lower extremity (Hamed et al., 2022; Hannah et al., 2017; Pal et al., 2007; Bujalski et al., 2018) and/or on variability introduced by measurement error (Myers et al., 2015). In contrast, the present study sought to determine: (1) the inherent variability of parameters used in modeling the elbow joint; (2) whether these parameters correlated to one another such that a subset of parameters could adequately predict other parameters; and (3) the parameters for which their variability has the most impact on musculoskeletal simulations (i.e., the parameters to which simulations are most sensitive). Our results indicate that although poorly correlated to one another, body parameters have greater relative variability than muscle parameters. However, it is muscle parameters that most influence inverse simulation results, meaning that even small errors in their approximation could have an outsized impact. These findings have implications for subject-specific modeling and for evaluating model robustness and accuracy. Both will be discussed below. Furthermore, there are several limitations to our study which should be acknowledged and will also be discussed.

### Subject-specific Modeling

Subject-specific models refer to models in which the parameter values are chosen to closely match measurements obtained from the individual of interest. This approach is preferred because it accounts for biological diversity, i.e. the model is customized per individual and measurement errors seem to have relatively small impact on simulation results (Myers et al., 2015; Valente et al., 2014; Hannah et al., 2017). However, obtaining these measurements is not practicable in all settings or circumstances. Measurements of muscle physiology, such as pennation angle or slack lengths require considerably sophisticated imaging techniques, such as MRI or ultrasound (Carbone et al., 2015; Scott et al., 1993; Parkkola et al., 1993; Hasson and Caldwell, 2012; Maganaris, 2001; O’Brien et al., 2010). These techniques, in turn, require appropriate expertise to collect, extract, and quantify these measurements. Therefore, the availability of the hardware and expertise needed to obtain these measurements limits the use of subject-specific models. In lieu of subject-specific measurements, parameter values may be estimated but these estimations require, by necessity, assumptions to be made. For example, parameter values may be linearly scaled to other anthropometric measurements, such as a segment’s length or a person’s weight (Winter, 2009). While these assumptions address the difficulties in creating subject-specific models, they may also introduce additional sources of error (Nolte et al., 2016). Here, we examined whether a subset of subject-specific parameters could be used to reasonably approximate other parameters. Unfortunately, we found generally weak correlations between parameters with two exceptions. Both muscle optimal length and pennation angle correlated well with the muscle’s maximum isometric force; although it is noteworthy that maximum isometric force had the fewest measurements in our dataset. Overall, our results confirm and expand upon previous work demonstrating the difficulty in approximating musculoskeletal parameters without direct measurements.

### Model Accuracy

The results of our study could be interpreted as evidence that generic or scaled models are insufficient or inaccurate. However, any criterion of model accuracy must consider the intended application. For example, some neuromechanical-based prosthetic controllers have exploited musculoskeletal models whose parameters are linearly scaled and/or empirically determined(Sartori et al., 2018). These models may not be accurate, in the strictest sense, to a specific individual but they are “good enough” for their particular application (Hicks et al., 2015). Here, we evaluated the impact of parameter variability independently of a specific application and found that muscle parameters generally have a larger impact on the optimized muscle activities generated by inverse simulations. Therefore, while a specific application will still require its own assessment of model accuracy, our results suggest that muscle parameters should be prioritized when it is determined that subject-specific parameters are needed.

### Limitations

Our study has some limitations that should be considered when interpreting or generalizing these results. First, our reduced elbow model was comprised of two muscle actuators with a single degree of freedom, while the human arm has many more of both. Although this simplified model has motor redundancy (2 control inputs for a single DOF), the larger number of muscle actuators in the human arm increases the potential solution space further, which may amplify the impact of musculoskeletal parameter variability on inverse simulation. Second, we constrained our Monte Carlo procedure to resample model parameters from a uniform distribution constrained by reported anthropometric measurements. These parameters did not include muscle geometry, which has recently been shown to be a significant source of variability in simulated ground reaction forces for the lower extremity (Hamed et al., 2022).

## CONCLUSIONS

In conclusion, musculoskeletal models have a wide range of potential applications, including performance assessment, orthosis/prosthesis design, and inferring neural control strategies. Different applications may have specific requirements on the accuracy of musculoskeletal simulations, i.e., some results may be “good enough” for one application and insufficient for another (Hicks et al., 2015). Our results may help inform future model development and applications. Our results demonstrate that models should prioritize approximating muscle parameters as accurately as possible to minimize simulation error, while body parameters may be sufficiently represented using mean values. Future work may: (1) investigate the range of parameter sensitivity to further constrain modeling assumptions; (2) determine whether parameter sensitivity scales linearly with model complexity; and (3) further disentangle sources of variability.

## AUTHOR CONTRIBUTIONS

Conceptualization/Study Design: RH

Data Collection: RH, BJ

Data Analysis: RH, BJ, DG

Writing: RH, DG

## ACKNOWLEDGEMENTS

This material is based upon work supported by the Department of Veterans Affairs, Veterans Health Administration, Office of Research and Development. We are grateful to Drs. Jonathan Wolpaw, Jonathan Carp, and Valeriya Gritsenko for their constructive feedback on the manuscript. The opinions expressed in this paper are those of the authors and do not represent the views of the Department of Veterans Affairs or the US Government.

