## Supplemental Data for "Variability and Impact of Musculoskeletal Modeling Parameters for the Human Elbow"

### Supplemental Material

#### 1 Additional Regression Results

##### 1.1 Full set of regression parameter estimates for main effects models

All regression parameter estimates of the bi error regression model are in Table 1, and for the tri error regression model are in Table 2. These tables complete corresponding Tables 9 and 10.

##### 1.2 Regression models with wo-way interactions

The top 50 regression model parameter estimates – ranked by magnitude of the estimate – for the regression model including two-way interactions with RMSE for bi error as the response are in Table 3. The corresponding values for tri error are in Table 4. The $r^2$ values are $r^2 = 0.89$ for the bi error model and $r^2 = 0.79$ for the tri error model, offering modest improvements over the corresponding $r^2$ values of $r^2 = 0.70$ and $r^2 = 0.65$ from the main effects models presented in the manuscript. This suggests that the terms that do have significant interactions, denoted with a colon (:) in Tables 3 and 4, have a meaningful effect on the RMSE.

Note in both tables the largest interactions are between the parameters such as muscle length and optimal tendon length that have the greatest effect on RMSE in the main effects models. This suggests that the effect of changes to one musculoskeletal parameter is dependent on the value of other musculoskeletal parameters. A more detailed tutoral on interpreting interaction effects is in the next section.

##### 1.3 Interpretation of two-way interactions

Since the data are mean-centered and scaled, interaction effects can be interpreted in general as follows using a simple example with two continuous predictor variables and a two-way interaction. Let $y$ be the observed standardized response variable (RMSE) and $x_1$ and $x_2$ be two observed musculoskeletal parameter values. The model with a two-way interaction between $x_1$ and $x_2$ would be expressed as

$$y = \beta_0 + \beta_1 x_1 + \beta_2 x_2 + \beta_{12} x_1 x_2$$

When $x_1$ is set to its mean value (zero) we get $y = \beta_0 + \beta_2 x_2$. When $x_1$ is set to, for example, one standard deviation above its mean value (so $x_1 = 1$) we get

$$y = \beta_0 + \beta_1 + \beta_2 x_2 + \beta_{12} x_2 = \beta_0 + \beta_1 + (\beta_2 + \beta_{12}) x_2$$

So as $x_1$ changes the effect on $x_2$ is an additive change in slope and intercept. For example, if $\beta_2 = 2$ and $\beta_{12} = 3$ the slope on $x_2$ increases from 2 to 5, thus leading to a larger increase in $y$ with increasing $x_2$. If $\beta_2 = 2$ and $\beta_{12} = -1$ the slope on $x_2$ is 1, resulting in a lesser increase in $y$ as $x_2$ increases. If $\beta_2 = 2$ and $\beta_{12} = -3$ the slope in $x_2$ changes sign, thus at mean $x_1$ $x_2$ has a positive slope and at $x_1 = 1$ $x_2$ has a negative slope.

| | Estimate | Std. Error | t value | P-value |
|---:|---:|---:|---:|---:|
| bi_maxforce | -0.45 | 0.02 | -25.55 | 0.00 |
| bi_tendonlen | 0.42 | 0.02 | 23.96 | 0.00 |
| tri_tendonlen | -0.36 | 0.02 | -20.19 | 0.00 |
| bi_optlen | 0.32 | 0.02 | 17.78 | 0.00 |
| tri_optlen | -0.21 | 0.02 | -11.55 | 0.00 |
| lower_mass | 0.19 | 0.02 | 10.41 | 0.00 |
| tri_maxforce | 0.17 | 0.02 | 9.70 | 0.00 |
| lower_comy | -0.09 | 0.02 | -5.01 | 0.00 |
| tri_penangle | 0.05 | 0.02 | 2.98 | 0.00 |
| lower_comx | -0.04 | 0.02 | -2.21 | 0.03 |
| upper_Iyz | 0.04 | 0.02 | 2.10 | 0.04 |
| upper_comz | 0.03 | 0.02 | 1.52 | 0.13 |
| upper_Ixx | 0.02 | 0.02 | 1.36 | 0.17 |
| lower_Izz | -0.02 | 0.02 | -1.23 | 0.22 |
| lower_Iyy | 0.02 | 0.02 | 1.13 | 0.26 |
| upper_Ixy | 0.02 | 0.02 | 1.12 | 0.26 |
| lower_Ixy | 0.01 | 0.02 | 0.83 | 0.41 |
| lower_Ixx | 0.01 | 0.02 | 0.66 | 0.51 |
| upper_comx | -0.01 | 0.02 | -0.63 | 0.53 |
| lower_comz | -0.01 | 0.02 | -0.51 | 0.61 |
| lower_Iyz | -0.01 | 0.02 | -0.49 | 0.62 |
| upper_Iyy | -0.01 | 0.02 | -0.46 | 0.65 |
| upper_mass | 0.00 | 0.02 | 0.24 | 0.81 |
| upper_Ixz | 0.00 | 0.02 | 0.23 | 0.82 |
| upper_comy | -0.00 | 0.02 | -0.20 | 0.84 |
| bi_penangle | -0.00 | 0.02 | -0.17 | 0.86 |
| upper_Izz | -0.00 | 0.02 | -0.02 | 0.98 |
| lower_Ixz | -0.00 | 0.02 | -0.01 | 0.99 |
| (Intercept) | 0.00 | 0.02 | 0.00 | 1.00 |

Table 1: Regression parameter estimates for bi error, main effects, all values.

| | Estimate | Std. Error | t value | P-value |
|---|---|---|---|---|
| tri_tendonlen | -0.45 | 0.02 | -23.63 | 0.00 |
| bi_optlen | -0.44 | 0.02 | -22.93 | 0.00 |
| bi_tendonlen | -0.39 | 0.02 | -20.30 | 0.00 |
| tri_penangle | 0.23 | 0.02 | 11.91 | 0.00 |
| tri_optlen | -0.09 | 0.02 | -4.52 | 0.00 |
| bi_maxforce | 0.09 | 0.02 | 4.51 | 0.00 |
| tri_maxforce | -0.05 | 0.02 | -2.44 | 0.01 |
| lower_comz | 0.04 | 0.02 | 2.13 | 0.03 |
| upper_mass | 0.04 | 0.02 | 1.99 | 0.05 |
| upper_comx | 0.04 | 0.02 | 1.90 | 0.06 |
| lower_comx | -0.04 | 0.02 | -1.85 | 0.07 |
| lower_Iyy | -0.03 | 0.02 | -1.82 | 0.07 |
| upper_Iyz | -0.03 | 0.02 | -1.74 | 0.08 |
| lower_Ixz | -0.03 | 0.02 | -1.72 | 0.09 |
| lower_Ixx | -0.03 | 0.02 | -1.58 | 0.11 |
| lower_Ixy | -0.03 | 0.02 | -1.51 | 0.13 |
| bi_penangle | 0.03 | 0.02 | 1.49 | 0.14 |
| upper_Ixz | -0.03 | 0.02 | -1.45 | 0.15 |
| lower_Izz | 0.02 | 0.02 | 0.87 | 0.39 |
| upper_Iyy | 0.02 | 0.02 | 0.84 | 0.40 |
| upper_Ixx | -0.01 | 0.02 | -0.53 | 0.60 |
| lower_Iyz | 0.01 | 0.02 | 0.49 | 0.62 |
| lower_comy | 0.01 | 0.02 | 0.40 | 0.69 |
| upper_Izz | -0.01 | 0.02 | -0.38 | 0.70 |
| upper_Ixy | 0.01 | 0.02 | 0.35 | 0.72 |
| upper_comz | 0.00 | 0.02 | 0.23 | 0.82 |
| upper_comy | 0.00 | 0.02 | 0.05 | 0.96 |
| lower_mass | 0.00 | 0.02 | 0.03 | 0.97 |
| (Intercept) | 0.00 | 0.02 | 0.00 | 1.00 |

Table 2: Regression parameter estimates for tri error, main effects, all values.

| | Estimate | Std. Error | t value | P-value |
|---:|---:|---:|---:|---:|
| bi_maxforce | -0.47 | 0.01 | -35.22 | 0.00 |
| bi_tendonlen | 0.40 | 0.01 | 29.79 | 0.00 |
| tri_tendonlen | -0.36 | 0.01 | -27.09 | 0.00 |
| bi_optlen | 0.33 | 0.01 | 24.95 | 0.00 |
| lower_mass | 0.20 | 0.01 | 15.16 | 0.00 |
| tri_optlen | -0.20 | 0.01 | -14.85 | 0.00 |
| bi_maxforce:bi_tendonlen | -0.19 | 0.01 | -14.27 | 0.00 |
| bi_maxforce:bi_optlen | -0.18 | 0.01 | -13.63 | 0.00 |
| tri_maxforce | 0.17 | 0.01 | 12.64 | 0.00 |
| tri_tendonlen:bi_tendonlen | -0.16 | 0.01 | -11.35 | 0.00 |
| bi_optlen:bi_tendonlen | 0.15 | 0.01 | 11.24 | 0.00 |
| tri_tendonlen:bi_optlen | -0.14 | 0.01 | -10.47 | 0.00 |
| tri_optlen:bi_tendonlen | -0.12 | 0.01 | -8.56 | 0.00 |
| tri_tendonlen:bi_maxforce | 0.10 | 0.01 | 7.48 | 0.00 |
| tri_maxforce:bi_tendonlen | 0.10 | 0.01 | 6.75 | 0.00 |
| lower_mass:bi_tendonlen | 0.09 | 0.01 | 6.11 | 0.00 |
| tri_maxforce:tri_tendonlen | -0.08 | 0.01 | -5.72 | 0.00 |
| tri_optlen:bi_maxforce | 0.07 | 0.01 | 4.80 | 0.00 |
| lower_comx | -0.06 | 0.01 | -4.70 | 0.00 |
| tri_optlen:bi_optlen | -0.07 | 0.01 | -4.64 | 0.00 |
| lower_comy | -0.05 | 0.01 | -3.93 | 0.00 |
| tri_maxforce:bi_optlen | 0.05 | 0.01 | 3.84 | 0.00 |
| tri_maxforce:bi_maxforce | -0.05 | 0.01 | -3.79 | 0.00 |
| lower_mass:bi_optlen | 0.05 | 0.01 | 3.77 | 0.00 |
| tri_optlen:tri_tendonlen | 0.05 | 0.01 | 3.63 | 0.00 |
| lower_mass:bi_maxforce | -0.05 | 0.01 | -3.42 | 0.00 |
| tri_maxforce:tri_optlen | -0.05 | 0.01 | -3.31 | 0.00 |
| lower_comy:bi_tendonlen | -0.04 | 0.01 | -2.92 | 0.00 |
| upper_mass:bi_penangle | -0.04 | 0.01 | -2.91 | 0.00 |
| upper_Ixx:upper_Iyy | 0.04 | 0.01 | 2.82 | 0.00 |
| upper_comx:upper_Ixx | 0.04 | 0.01 | 2.79 | 0.01 |
| lower_comz:lower_Iyz | 0.04 | 0.01 | 2.54 | 0.01 |
| upper_comz:tri_tendonlen | -0.03 | 0.01 | -2.49 | 0.01 |
| upper_Ixz:tri_maxforce | 0.03 | 0.01 | 2.34 | 0.02 |
| lower_Ixy:tri_maxforce | -0.03 | 0.01 | -2.32 | 0.02 |
| lower_comy:bi_optlen | -0.03 | 0.01 | -2.31 | 0.02 |
| upper_Ixx:lower_Ixz | 0.03 | 0.01 | 2.30 | 0.02 |
| lower_Iyy:bi_tendonlen | 0.03 | 0.01 | 2.23 | 0.03 |
| upper_Izz:lower_Ixz | 0.03 | 0.01 | 2.22 | 0.03 |
| upper_comx:upper_comy | -0.03 | 0.01 | -2.20 | 0.03 |
| upper_Ixz:bi_tendonlen | -0.03 | 0.01 | -2.20 | 0.03 |
| lower_comz:lower_Ixz | 0.03 | 0.01 | 2.18 | 0.03 |
| upper_Ixx:upper_Izz | 0.03 | 0.01 | 2.18 | 0.03 |
| lower_mass:lower_Ixx | 0.03 | 0.01 | 2.16 | 0.03 |
| lower_comz:tri_penangle | -0.03 | 0.01 | -2.15 | 0.03 |
| tri_penangle:bi_maxforce | -0.03 | 0.01 | -2.11 | 0.04 |
| lower_mass:bi_penangle | -0.03 | 0.01 | -2.06 | 0.04 |
| upper_Ixz:lower_comy | 0.03 | 0.01 | 2.03 | 0.04 |
| upper_Ixz:lower_Izz | 0.03 | 0.01 | 1.99 | 0.05 |
| lower_comz:lower_Ixy | -0.03 | 0.01 | -1.99 | 0.05 |

Table 3: Regression parameter estimates for bi error, two-way interactions, top 50 by p value.

| | Estimate | Std. Error | t value | P-value |
|---|---|---|---|---|
| tri_tendonlen | -0.47 | 0.02 | -25.43 | 0.00 |
| bi_optlen | -0.42 | 0.02 | -22.98 | 0.00 |
| bi_tendonlen | -0.39 | 0.02 | -20.58 | 0.00 |
| tri_penangle | 0.22 | 0.02 | 11.89 | 0.00 |
| tri_tendonlen:bi_optlen | 0.21 | 0.02 | 11.12 | 0.00 |
| tri_tendonlen:bi_tendonlen | 0.22 | 0.02 | 11.04 | 0.00 |
| bi_optlen:bi_tendonlen | 0.15 | 0.02 | 7.77 | 0.00 |
| tri_penangle:bi_optlen | -0.12 | 0.02 | -6.47 | 0.00 |
| tri_penangle:bi_tendonlen | -0.10 | 0.02 | -5.33 | 0.00 |
| bi_maxforce | 0.07 | 0.02 | 3.84 | 0.00 |
| tri_optlen | -0.06 | 0.02 | -3.43 | 0.00 |
| lower_comy:tri_maxforce | -0.07 | 0.02 | -3.32 | 0.00 |
| tri_maxforce:tri_tendonlen | 0.06 | 0.02 | 3.27 | 0.00 |
| tri_optlen:bi_tendonlen | 0.06 | 0.02 | 3.25 | 0.00 |
| lower_comx:tri_tendonlen | 0.06 | 0.02 | 3.19 | 0.00 |
| upper_comy:bi_tendonlen | 0.05 | 0.02 | 2.81 | 0.01 |
| tri_optlen:bi_optlen | 0.05 | 0.02 | 2.78 | 0.01 |
| lower_comx:bi_maxforce | -0.05 | 0.02 | -2.61 | 0.01 |
| upper_Ixx:upper_Ixz | 0.05 | 0.02 | 2.48 | 0.01 |
| upper_Izz:tri_maxforce | 0.05 | 0.02 | 2.44 | 0.02 |
| lower_comy:bi_penangle | 0.05 | 0.02 | 2.43 | 0.02 |
| upper_Iyy:lower_Ixx | -0.05 | 0.02 | -2.35 | 0.02 |
| upper_comy:tri_penangle | 0.05 | 0.02 | 2.34 | 0.02 |
| upper_Ixy:tri_penangle | 0.04 | 0.02 | 2.33 | 0.02 |
| upper_mass:upper_comx | 0.04 | 0.02 | 2.30 | 0.02 |
| upper_Ixx:bi_penangle | -0.04 | 0.02 | -2.29 | 0.02 |
| lower_comz:lower_Ixz | -0.04 | 0.02 | -2.28 | 0.02 |
| lower_Ixx:tri_penangle | -0.04 | 0.02 | -2.27 | 0.02 |
| upper_Ixx:lower_Iyy | 0.04 | 0.02 | 2.24 | 0.03 |
| lower_comx | -0.04 | 0.02 | -2.23 | 0.03 |
| lower_comy:bi_maxforce | -0.04 | 0.02 | -2.20 | 0.03 |
| lower_Iyy:bi_optlen | 0.04 | 0.02 | 2.17 | 0.03 |
| upper_Iyz:lower_Ixz | -0.04 | 0.02 | -2.14 | 0.03 |
| upper_Iyz:lower_mass | 0.04 | 0.02 | 2.13 | 0.03 |
| lower_Ixx:lower_Ixz | 0.04 | 0.02 | 2.12 | 0.03 |
| upper_mass:tri_optlen | -0.04 | 0.02 | -2.10 | 0.04 |
| bi_maxforce:bi_optlen | -0.04 | 0.02 | -2.10 | 0.04 |
| upper_Iyz:lower_comy | -0.04 | 0.02 | -2.09 | 0.04 |
| upper_comx:tri_penangle | 0.04 | 0.02 | 2.09 | 0.04 |
| upper_Ixx:bi_maxforce | -0.04 | 0.02 | -2.08 | 0.04 |
| upper_comx:lower_Ixy | -0.04 | 0.02 | -2.07 | 0.04 |
| upper_comx:upper_Ixz | 0.04 | 0.02 | 2.06 | 0.04 |
| upper_Ixz:lower_comz | -0.04 | 0.02 | -2.04 | 0.04 |
| upper_Ixy:lower_comx | -0.04 | 0.02 | -2.03 | 0.04 |
| upper_mass | 0.04 | 0.02 | 2.03 | 0.04 |
| upper_Izz:upper_Ixy | 0.04 | 0.02 | 2.03 | 0.04 |
| upper_Izz:bi_optlen | 0.04 | 0.02 | 2.01 | 0.04 |
| upper_mass:tri_maxforce | 0.04 | 0.02 | 2.00 | 0.05 |
| upper_Ixy:lower_Iyy | -0.04 | 0.02 | -2.00 | 0.05 |
| lower_comy:lower_comz | 0.04 | 0.02 | 1.97 | 0.05 |

Table 4: Regression parameter estimates for tri error, two-way interactions. Top 50 by p value.
